# Kinetic And Structural Parameters Governing Fic-Mediated Adenylylation/AMPylation of the Hsp70 chaperone, BiP/GRP78

**DOI:** 10.1101/494930

**Authors:** Anwesha Sanyal, Erica A. Zbornik, Ben G. Watson, Charles Christoffer, Jia Ma, Daisuke Kihara, Seema Mattoo

## Abstract

Fic (filamentation induced by cAMP) proteins regulate diverse cell signaling events by post-translationally modifying their protein targets, predominantly by the addition of an AMP (adenosine monophosphate). This modification is called Fic-mediated Adenylylation or AMPylation. We previously reported that the human Fic protein, HYPE/FicD, is a novel regulator of the unfolded protein response (UPR) that maintains homeostasis in the endoplasmic reticulum (ER) in response to stress from misfolded proteins. Specifically, HYPE regulates UPR by adenylylating the ER chaperone, BiP/GRP78, which serves as a sentinel for UPR activation. Maintaining ER homeostasis is critical for determining cell fate, thus highlighting the importance of the HYPE-BiP interaction. Here, we study the kinetic and structural parameters that determine the HYPE-BiP interaction. By measuring the binding and kinetic efficiencies of HYPE in its activated (Adenylylation-competent) and wild type (de-AMPylation-competent) forms for BiP in its wild type and ATP-bound conformations, we determine that HYPE displays a nearly identical preference for the wild type and ATP-bound forms of BiP *in vitro* and preferentially de-AMPylates the wild type form of adenylylated BiP. We also show that AMPylation at BiP’s Thr366 versus Thr518 sites differentially affect its ATPase activity, and that HYPE does not adenylylate UPR accessory proteins like J-protein ERdJ6. Using molecular docking models, we explain how HYPE is able to adenylylate Thr366 and Thr518 sites *in vitro.* While a physiological role for AMPylation at both the Thr366 and Thr518 sites has been reported, our molecular docking model supports Thr518 as the structurally preferred modification site. This is the first such analysis of the HYPE-BiP interaction and offers critical insights into substrate specificity and target recognition.

## Introduction

Fic (Filamentation induced by cyclic AMP) proteins are a recently discovered class of enzymes that carry out diverse post-translational modifications (1,2). Predominant amongst these is a modification called Adenylylation or AMPylation, which entails the covalent addition of an AMP to the target protein. Adenylylation/AMPylation occurs on Thr, Tyr, or Ser residues of the target protein. The AMPylation event most resembles the adenylylation reaction described by Earl Stadtman in the 1960s, where *E. coli* glutamine synthetase (GS) is reversibly adenylylated by GS adenylyltransferase (GS-ATase), and that carried out by the newly discovered Sel-O atypical kinase (3,4). Despite the similarity of these reactions, Fic proteins share no sequence similarity with GS-ATase or any other known adenylyltransferases (5). Instead, they are defined by a Fic domain, which consists of six to eight α-helices surrounding a conserved HxFx(D/E)(G/A)N(G/K)RXXR motif (6,7). The invariant His residue of this Fic motif is required for catalytic activity. Further, many Fic proteins are intrinsically regulated by an inhibitory helix (α_inh_) defined by an (S/T)xxxE(G/N) motif that blocks nucleotide docking in the Fic active site (8). Structurally, the catalytic loop positions the α-phosphate of the nucleotide for a nucleophilic attack by the target Tyr, Thr or Ser hydroxyl moiety, while the conserved His of the Fic active site functions as a general base to deprotonate the nucleophile (6,8). Additionally, some Fic proteins utilize ATP to phosphorylate their target proteins, while still others show specificity for UTP or CDP-choline to add a UMP or phosphocholine, respectively, to their targets (5,9-13).

The first reports of Fic-mediated adenylylation were described for bacteria. Specifically, *Vibrio parahemolyticus* and *Histophilus somni* were shown to secrete Fic-domain-containing bacterial effectors -VopS and IbpA, respectively - which induced toxicity in host cells by adenylylating small GTPases, RhoA, Rac1 and Cdc42 (5,10). A co-crystal structure of IbpA in complex with Cdc42 revealed that adenylylation requires IbpA to interact with two structural loops (Switch I and Switch II) on Cdc42, with adenylylation occurring only on the Switch I loop (6). Consequently, peptides representing just the Switch I region fail to be recognized or adenylylated by IbpA (1). Further, the IbpA-Cdc42 co-crystal structure showed that adenylylation locks Cdc42 in an inactive conformation (6). Extending these findings to VopS, Xiao *et al*. predicted and validated a similar model of target recognition (6). To date, the IbpA-Cdc42 structure offers the only snapshot of Fic-mediated adenylylation, target recognition, and ATP-binding in the context of both the Fic enzyme and its target. As new targets for Fic proteins become known, co-crystal structures of these complexes become imperative to understand how Fic proteins recognize and modify their targets.

Fic proteins are evolutionarily conserved. We characterized the sole human Fic protein, HYPE/FicD, as an adenylyltransferase that functions to regulate ER stress in mammalian cells (Figure 1A) (14). HYPE (Huntingtin yeast interacting protein E) was identified in a yeast-two hybrid screen to interact with the Huntingtin protein, mutations within which cause Huntington’s disease (15). The physiological relevance of this interaction remains unknown. HYPE is well conserved in eukaryotes, displaying significant amino acid sequence similarity to homologs in *M. musculus* (89.5%), *C. elegans* (45.4%), *D. rerio* (76%) and *D. melanogaster* (55%). HYPE is expressed in all tissues, albeit at very low levels (1,5). However, treatment of HEK293 human kidney epithelial cells with drugs that induce ER stress induces an up-regulation of HYPE expression (14). Additionally, an E234G mutation in HYPE’s α-inhibitory helix renders a constitutively active adenylyltransferase (8,14). Consequently, constitutive expression of adenylylation-competent HYPE by transfecting HEK293 cells with E234G-HYPE induces apoptosis, highlighting the importance of maintaining tight regulation of HYPE’s enzymatic activity in the cells (14).

**Figure 1:**
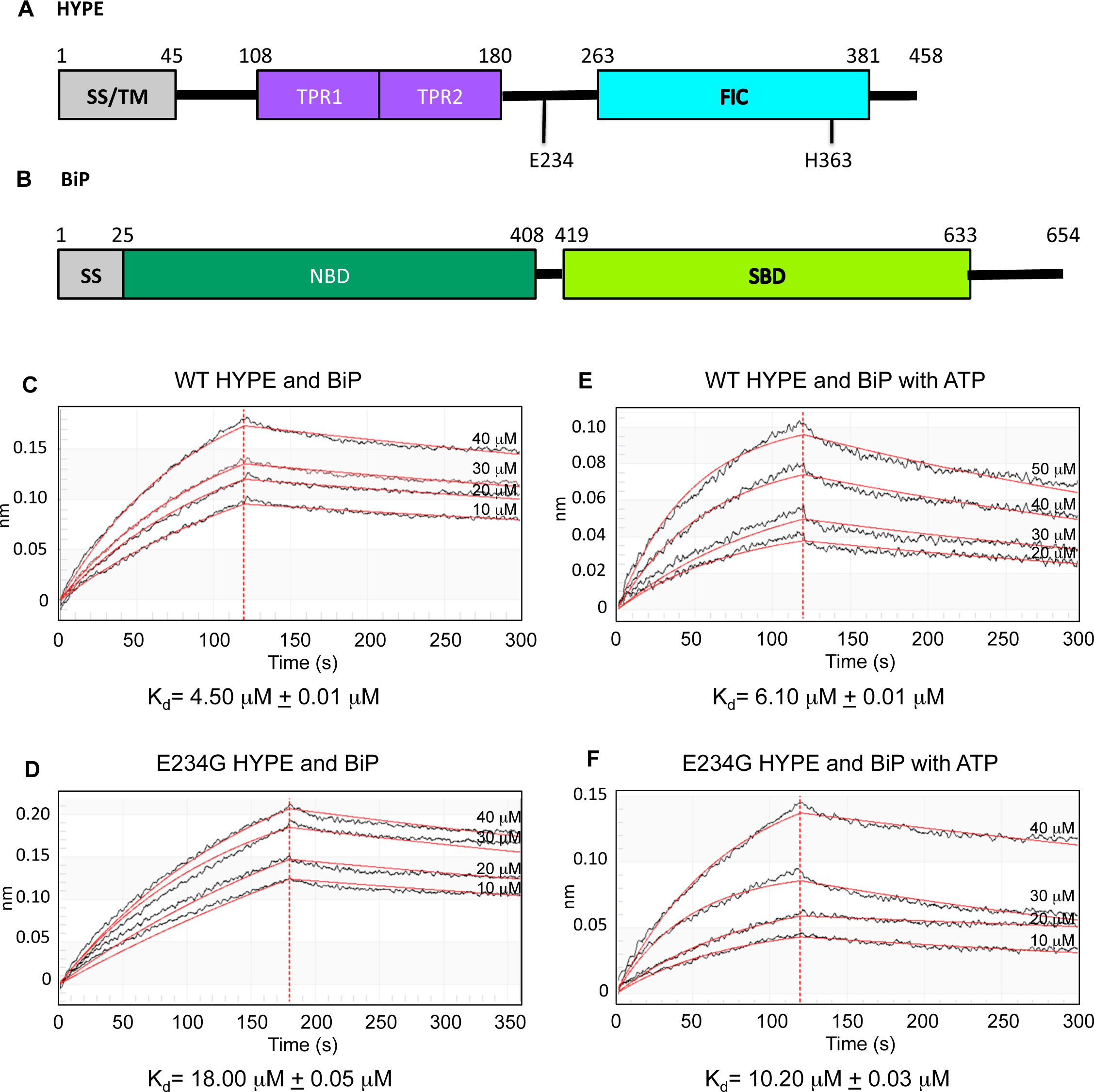
A) Schematic representation of HYPE. Residues 1-45 represent the signal sequence (SS) and hydrophobic, predicted transmembrane (TM) domain of HYPE. HYPE’s two tetratricopeptide regions (TPR1 and TPR2) precede the α_inh_ region, which contains the E234 regulatory residue. Mutation of E234G renders HYPE a constitutively active adenylyltransferase. The catalytic His363 of HYPE’s Fic domain (cyan) is shown. Mutation of H363A renders HYPE catalytically inactive for both its adenylyltransferase and de-AMPylase activities. **B) Schematic representation of BiP.** Residues 1-25 represent a predicted ER signal sequence followed by the nucleotide-binding domain (NBD) which can bind ATP or ADP. A flexible linker connects BiP’s NBD (dark green) to its substrate-binding domain (SBD; light green), which binds misfolded client proteins. This flexible linker allows BiP to assume its “open” and “closed” conformations. **(C-F) WT- and E234G-HYPE bind BiP with similar affinities.** The steady state binding affinity for (C) WT-HYPE or (D) E234G-HYPE with BiP was obtained by biolayer interferometry using Octet Red from ForteBio. A constant concentration of BiP was immobilized on HIS1K sensors and incubated with varying concentrations of WT- or E234G-HYPE. Binding affinities were also measured in the presence of ATP for (E) WT-HYPE and BiP and (F) E234G-HYPE and BiP. The data were fitted to a 1:1 global fitting model.

We previously demonstrated that HYPE localizes to the lumen of the ER and adenylylates the ER Hsp70 chaperone, BiP (Binding to immunoglobulin protein; Figure 1B) (14). BiP is a key regulator of the ER homeostasis pathway called UPR (unfolded protein response) that relieves ER stress via three signal transducers in the ER membrane: IRE1, ATF6 and PERK (16). BiP consists of a nucleotide-binding domain (NBD) that binds and hydrolyzes ATP and a substrate-binding domain (SBD) that binds misfolded proteins (17,18). BiP’s ATPase activity is required for it to refold misfolded proteins (19). We reported that HYPE catalyzes a single adenylylation on human BiP at Ser365 or Thr366 (14). Both these residues lie in BiP’s NBD. The Thr366 site is targeted for adenylylation by HYPE’s homolog in *Drosophila melanogaster* (dFic) as well, while *C. elegans* Fic (Fic-1) adenylylates *C. elegans* BiP at various residues in the NBD, including the T366 equivalent, T371 (20,21). Interestingly, we found adenylylation at Thr366 enhanced BiP’s ATPase activity *in vitro*, albeit by a modest two-fold increase over wild type BiP (14). Thereafter, a second adenylylation within BiP’s SBD at Thr518 was reported (22). This Thr518 modification was shown to inhibit BiP’s ATPase activity *in vitro*, again with a modest two-fold decrease compared to wild type BiP (22). Both the above ATPase assays were conducted *in vitro* using HYPE and BiP in isolation. However, the ATPase cycle of BiP involves the transfer of misfolded proteins from another set of Hsp40 chaperones called J proteins. In an elegant study, it was recently reported that adenylylation at Thr518 specifically inhibits BiP’s J-protein assisted ATPase activity (23). Based on assessment of BiP adenylylation within cells, it has been proposed that BiP is predominantly adenylylated at Thr518 (21), while another report describes a physiological role for adenylylation at Thr366 in neurodegeneration in a *D. melanogaster* model (24). The physiological signal(s) that activates HYPE’s enzymatic activity is unknown.

By monitoring protein misfolding within the ER lumen, BiP serves as a sentinel for activating the UPR. Thus, by virtue of its ability to adenylylate and alter BiP’s enzymatic activity, we established HYPE as a key player in maintaining ER homeostasis (14). Indeed, knockdown of HYPE prevents proper UPR progression and renders cells more susceptible to death under sustained ER stress (14). The absence of HYPE appears to specifically alter activation of the ATF6 and PERK branches of the UPR pathway, with negligible effect on the IRE1 branch (14). Indeed, it was recently shown that IRE1 can be activated independently of BiP activity or binding of misfolded proteins (25).

HYPE is a 52kDa protein that consists of a hydrophobic N-terminus, followed by two tetratricopeptide (TPR) repeats, and a canonical Fic domain (Figure 1A) (5). Additionally, between the second TPR domain and the Fic domain, HYPE has an α-inhibitor sequence (Figure 1A). As indicated earlier, mutation of the conserved Glu (E234G) in the α_inh_ of HYPE relieves its intrinsic inhibition and renders it constitutively active, while mutation of the conserved His (H363A) in the Fic active site renders it inactive. Though several lines of evidence assign a weak, basal adenylylation activity to WT-HYPE (1,5,14,21), Preissler *et al*. show that WT-HYPE predominantly catalyzes the removal of the AMP moiety from target proteins (26). Thus, E234G-HYPE functions as the active adenylyltransferase and WT-HYPE functions predominantly as a de-adenylylating enzyme or “de-AMPylase”. Together, these findings further highlight the stringent control cells exert on both HYPE expression and its enzymatic activity.

The crystal structure of HYPE reveals that it exists as a dimer (27). Two HYPE monomers are positioned in an antiparallel conformation such that both the Fic active site clefts and the TPR grooves are exposed for possible substrate binding and catalysis. Analysis of cofactor binding indicates that both WT- and activated E234G-HYPE have equally high propensity to bind to ADP, while only E234G-but not WT-HYPE binds ATP. A comprehensive study of the α_inh_ domain of Fic proteins suggested that a structural rearrangement may be required to activate HYPE (8). However, the overall structures of WT-and E234G-HYPE are similar, indicating that the inhibition is primarily due to charge repulsion of ATP caused by E234 and not due to a gross change in conformation (24).

Here, we conduct a detailed kinetic characterization of HYPE-mediated adenylylation and de-AMPylation of BiP. We determine the reaction rates for wild type versus activated E234G-HYPE, and find that even though E234G-HYPE is far more efficient at adenylylating BiP, both enzymes bind BiP with comparable affinities. We also assess the kinetics of BiP AMPylation by E234G-HYPE, as well as the ability of WT-HYPE to de-AMPylate BiP in its wild type and ATP-bound conformations. We assess the consequence of BiP AMPylation with respect to modification at its Ser365/Thr366 and Thr518 sites, alone and in the presence of J protein, ERdJ6. Our data show differential effects of adenylylation at the Thr366 versus Thr518 sites, and that J protein itself is not a target of HYPE-mediated adenylylation. We delineate the kinetics for HYPE-mediated adenylylation of BiP’s NBD versus SBD, and find that HYPE can modify BiP’s T366 in the absence of an SBD but needs both NBD and SBD in order to adenylylate Thr518 *in vitro*. Finally, building on the results of our kinetic analyses and applying them to the known crystal structures for HYPE and BiP, we computationally model the HYPE-BiP complex to reveal residues critical for interaction. Our molecular docking models explain how BiP’s NBD serves as an adenylylation target for HYPE *in vitro*, while modification at the SBD requires HYPE to interact with both BiP’s NBD and SBD. Further, our docking models show that despite the *in vitro* kinetic values that support adenylylation in BiP’s NBD, Thr518 in BiP’s SBD is structurally the more physiologically preferred site of adenylylation, in agreement with *in vivo* data from Preissler *et al.* (22). This is the first kinetic and structural characterization of the HYPE-BiP interaction and offers valuable insight for manipulating this fundamental interaction that governs cell fate in response to ER stress. Taken together, our structural docking and kinetic analyses explain the *in vitro* interactions between HYPE and BiP – an important first step towards the development of therapeutics and small molecules for manipulating this critical cell signaling nexus.

## Results

### Wild type and E234G-HYPE bind BiP with similar affinities

We previously characterized WT-HYPE as a weak adenylyltransferase, but its physiological target had remained elusive (5). Eventually, using the catalytically inactive H363A-HYPE mutant as a substrate trap, we successfully immunoprecipitated and identified BiP as an interacting partner for HYPE (14). Further, using the activated E234G-HYPE mutant, we established BiP as a physiologically relevant adenylylation target for HYPE. Indeed, we and others have now reported Thr366 and Thr518 as two sites of adenylylation for BiP (14,21,22). We, therefore, quantified the HYPE-BiP interaction by assessing their binding affinities using Biolayer Interferometry. Specifically, we immobilized bacterially expressed and purified BiP-His_6_ on a His-antibody sensor and titrated increasing concentrations of untagged WT- or E234G-HYPE. By measuring the rates of association and dissociation of HYPE, we determined the K_d_ of binding for WT-HYPE as 4.50 μM ± 0.01 μM and for E234G-HYPE as 18.00 μM ± 0.05 μM. Since these K_d_ values are within a micromolar range, we infer that WT and E234G HYPE bind BiP with similar binding affinities (Figures 1C and 1D). This observation is in accordance with the fact that the crystal structures of WT- and E234G-HYPE are not significantly different (27).

We next asked whether HYPE’s binding affinity for BiP changes in the presence of ATP. Addition of ATP did not alter the binding affinities of BiP for WT-HYPE (K_d_ = 6.10 μM ± 0.01 μM) or E234G-HYPE (K_d_ = 10.20 μM ± 0.03 μM), indicating that ATP is not a prerequisite for substrate binding and, instead, simply serves as a necessary component of the adenylylation reaction (Figures 1E and 1F). Further, the data for the binding curve fitted best to a 1:1 fitting model, suggesting that one molecule of HYPE binds to a single molecule of BiP. Since, HYPE crystallizes as a stable dimer (27), we propose that the HYPE-BiP complex most likely exists as a 2:2 complex consisting of a single HYPE dimer bound to two individual BiP molecules. Such an interaction of one Fic dimer and two substrate molecules is reminiscent of the bacterial Fic protein IbpA in complex with its Rho GTPase substrate, Cdc42 (6).

### Reaction kinetics of wild-type versus E234G-HYPE-mediated BiP adenylylation

*In vitro* adenylylation assays using α^32^P-ATP as a nucleotide source indicate that E234G-HYPE efficiently hydrolyzes ATP to AMP and PPi (inorganic pyrophosphate) to catalyze the addition of AMP to BiP (1). To determine the kinetic constants for HYPE-mediated adenylylation of BiP, we first performed time-dependent kinetics over a range of BiP concentrations to determine the linear range of this reaction for E234G-HYPE. Using WT-BiP (aa19-637) as the adenylylation target at concentrations ranging from 0.5-2 μM, the *in vitro* adenylylation reaction was initiated with a constant amount of the E234G-HYPE enzyme and constant α^32^P-ATP for a reaction time ranging from 0 to 45 minutes. As evident from Figures 2A, a reaction time of 4 minutes represents the adenylylation reaction in its linear range where 10% of BiP is adenylylated. Subsequent kinetic experiments were, therefore, performed at a reaction time of 4 minutes. For the determination of Km and kcat of the adenylylation reaction, *in vitro* adenylylation of BiP was conducted using a constant concentration of E234G-HYPE (0.1 μM) and ATP (1 mM) while the concentration of WT-BiP was increased from 0 μM to 15 μM. The apparent Km of E234G-HYPE for BiP was 2.37 ± 0.54 μM, and the kcat of the reaction catalyzed by E234G-HYPE was 0.524 ± 0.038 s^-1^ (Figure 2B). We also attempted to determine kinetic parameters for WT-HYPE incubated with WT-BiP or T229A-BiP (data not shown). However, as expected, these reactions did not reach saturation under steady state conditions to fit a true Michael-Menten curve because WT-HYPE displays simultaneous weak adenylylating and stronger de-AMPylating activities.

**Figure 2:**
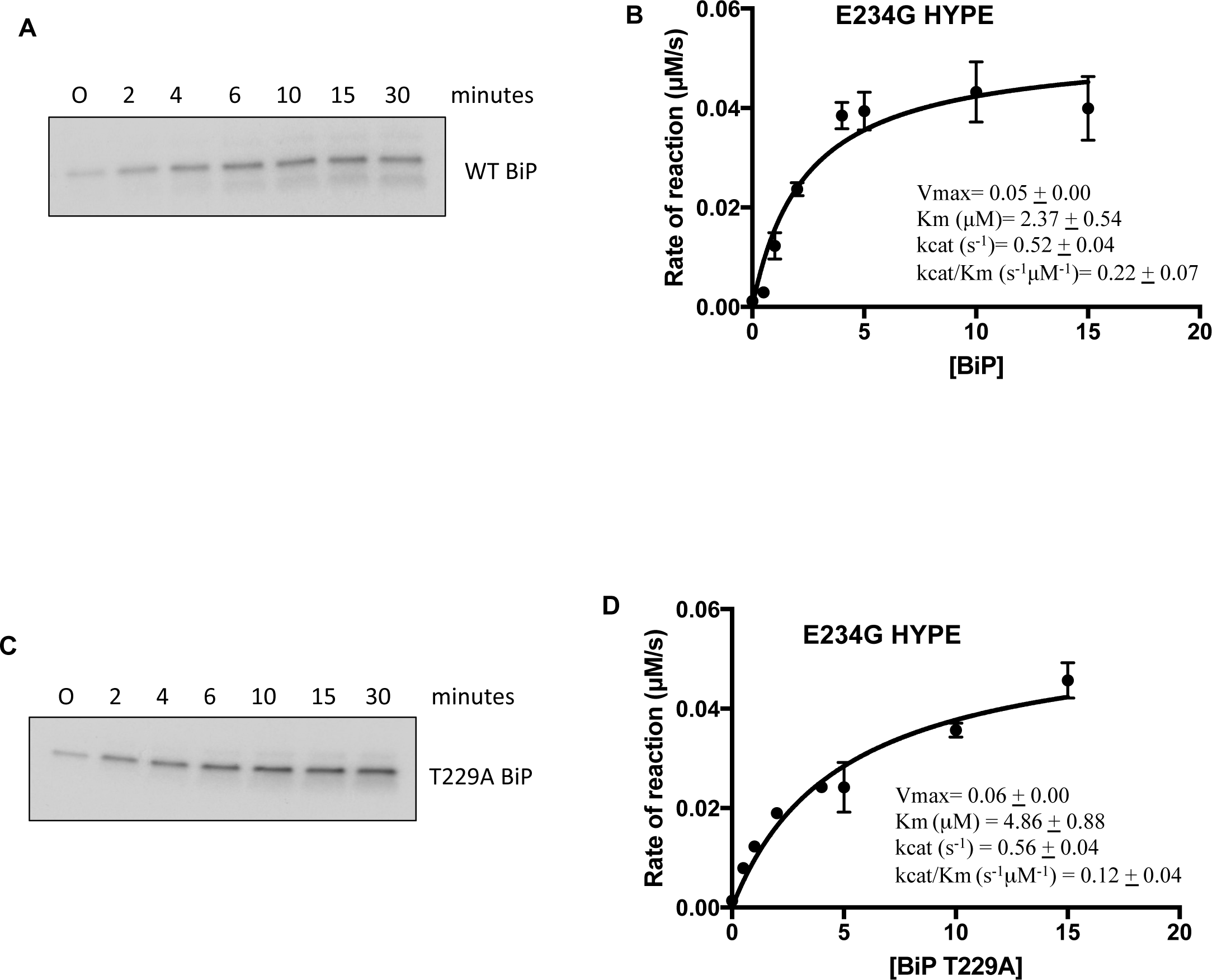
Steady-state kinetic analysis of WT- or E234G-HYPE for BiP and T229A-BiP. Linear range of the adenylylation reaction was determined using E234G-HYPE and (A) BiP or (C) T229A-BiP over a range of BiP concentrations (0.5, 1, 1.5, and 2 μM) at different time points from 0-30 minutes. A representative autoradiograph with 1 μM WT- or T229A-BiP is shown (A and C, respectively). Kinetic parameters for WT-BiP or T229A-BiP were measured using constant concentrations of ATP and enzyme (E234G-HYPE), while varying the concentrations of substrate BiP (B) or T229A-BiP (D). Assays were performed in triplicates and data was fitted to Michaelis-Menten equation in Graph-Pad Prism. Error bars indicate S.D.

### In vitro adenylylation kinetics do not support HYPE’s preference for the ATP-bound form of BiP as a substrate

BiP is an Hsp70 molecular chaperone that hydrolyzes ATP and utilizes the resultant energy for folding unfolded or misfolded proteins (17). In the cell, BiP exists in two conformations: 1) an ADP-bound “open” conformation that binds its misfolded protein substrates with high affinity, and 2) an ATP-bound “closed” conformation that folds and releases its protein substrates at a faster rate. Recently, it was suggested that the ATP-bound form of BiP is a better substrate for HYPE-mediated adenylylation (22). We, therefore, tested this assertion by comparing the reaction kinetics for HYPE’s ability to adenylylate WT-BiP versus a Thr229 to Ala mutant of BiP, which mimics its ATP-bound conformation. T229A-BiP is a well-characterized mutant of BiP that is unable to hydrolyze ATP and hence remains locked in the ATP-bound state (18). As described in the Methods, using an *in vitro* adenylylation reaction with α^32^P-ATP as a nucleotide source and increasing concentrations of WT or T229A-BiP, we determined that the linear range for the adenylylation of T229A-BiP matches that for WT-BiP (compare Figures 2A to 2C). Therefore, adenylylation kinetics were assessed at a reaction time of 4 minutes. The kinetic parameters of adenylylation-competent E234G-HYPE for T229A-BiP (ie., Km = 4.86 μM ± 0.88 μM and kcat = 0.56 s^-1^ ± 0.04 s^-1^) match those for WT-BiP (ie., Km = 2.37 μM ± 0.54 μM and kcat = 0.524 s^-1^ ± 0.038 s^-1^) (Figures 2B and 2D). Additionally, the efficiency of reaction by E234G-HYPE for T229A-BiP as measured by kcat/Km = 0.115 s^-1^μM^-1^ is only 2-fold lower than for WT-BiP (kcat/Km = 0.22 ± 0.07 s^-1^μM^-1^). These data suggest that WT-and ATP-bound forms of BiP are adenylylated by E234G-HYPE with similar efficiency. Accordingly, steady state binding assays using either WT- or E234G-HYPE indicate that both forms of HYPE bind to T229A-BiP with similar affinity, ranging from K_d_ = 2.5-2.8 μM (Figures 3A and 3B). Addition of ATP does not significantly alter the binding (Figures 3C and 3D).

**Figure 3:**
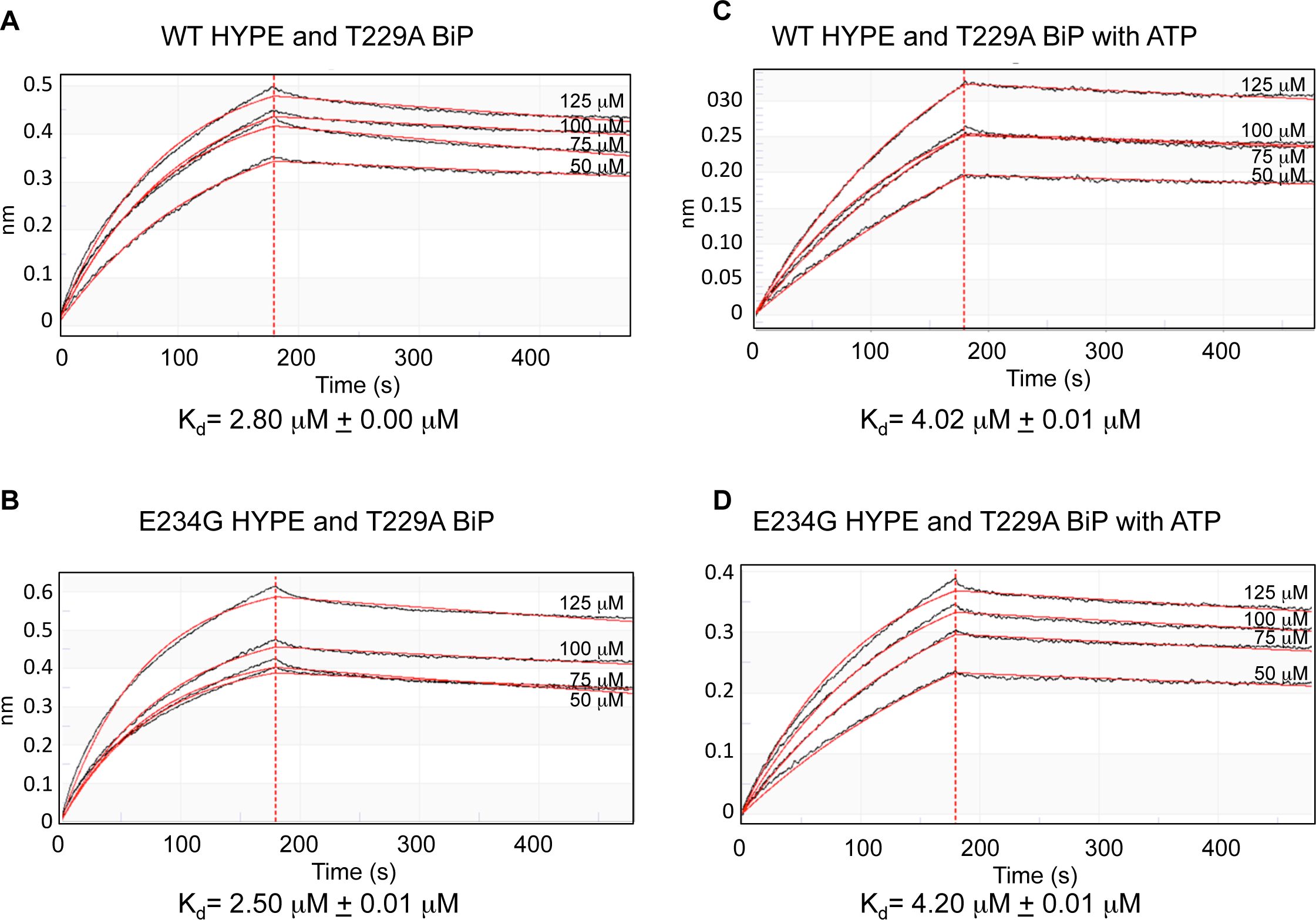
WT- and E234G-HYPE bind T229A-BiP with similar affinities. Steady state binding affinity determinations for (A) WT-HYPE or (B) E234G-HYPE incubated with T229A-BiP-His_6_ immobilized on HIS1K sensors were obtained by Biolayer Interferometry. Similar binding affinities were also observed in the presence of ATP for (C) WT-HYPE and T229A-BiP and (D) E234G-HYPE and T229A-BiP.

Overall, while our adenylylation data do corroborate a previous report that states that ATP-bound BiP is preferentially adenylylated by E234G-HYPE (compare Km/Kcat values for Figures 2B versus 2D), these differences are not significant at least in an *in vitro* experimental scenario. It remains to be determined whether ATP-bound BiP is the preferred adenylylation target *in vivo*, especially in the context of a misfolded protein client.

### Kinetic parameters for the de-AMPylation of BiP

We previously showed that Fic-mediated adenylylation could be reversed by treatment with phosphodiesterases (5). Indeed, bacteria have been shown to encode Fic proteins in operons coupled with the reversing enzyme (28). Further support for such reversible adenylylation comes from the adenylylation of GS by GS-ATase in *E. coli*, where the GS-ATase itself can add or remove AMP in response to signal-specific activation (29). Adenylylation of BiP is also reported to be transient, suggesting that enzyme(s) capable of removing the AMP from BiP must exist in cells (20). It is hypothesized that reversible adenylyation of BiP is a means of tightly controlling UPR. Recently, it was discovered that like GS-ATase, WT-HYPE itself catalyzes the de-adenylylation/de-AMPylation of BiP (29). Specifically, it is proposed that during the de-AMPylation reaction, residue E234 in WT-HYPE coordinates a water molecule, which attacks the phosphodiester bond between BiP and AMP, thereby removing the AMP. Accordingly, mutation of E234 switches the enzyme from a de-AMPylase to an adenylyltransferase, presumably by removing the charge repulsion between E234 and the α-phosphate of ATP. Thus, we sought to test the de-AMPylation activity of WT-HYPE by incubating adenylylated BiP (BiP-AMP) with increasing concentrations of WT-HYPE to determine the kinetics of de-AMPylation. For this, we first adenylylated BiP by incubating WT-BiP with E234G-HYPE in the presence of α^32^P-ATP until the adenylylation reaction reached saturation. Next, the sample was left untreated or incubated for 5 to 50 minutes with a constant amount of WT-HYPE to determine the linear range of the de-AMPylation reaction (Figure 4A). Our data confirm that WT-HYPE functions as a de-AMPylase (Figures 4A and 4B). Specifically, incubation with WT-HYPE for 50 minutes represented conditions where over 80% of the radiolabel was removed from BiP-AMP (Figure 4B). Interestingly, E234G-HYPE displayed only a 50% loss of the radiolabel at this time point, which may be due to simultaneous autoadenylylation. From this reaction, we calculated the linear range to be at 30 minutes, when approximately 50% of the de-AMPylation reaction is catalyzed for BiP and HYPE.

**Figure 4:**
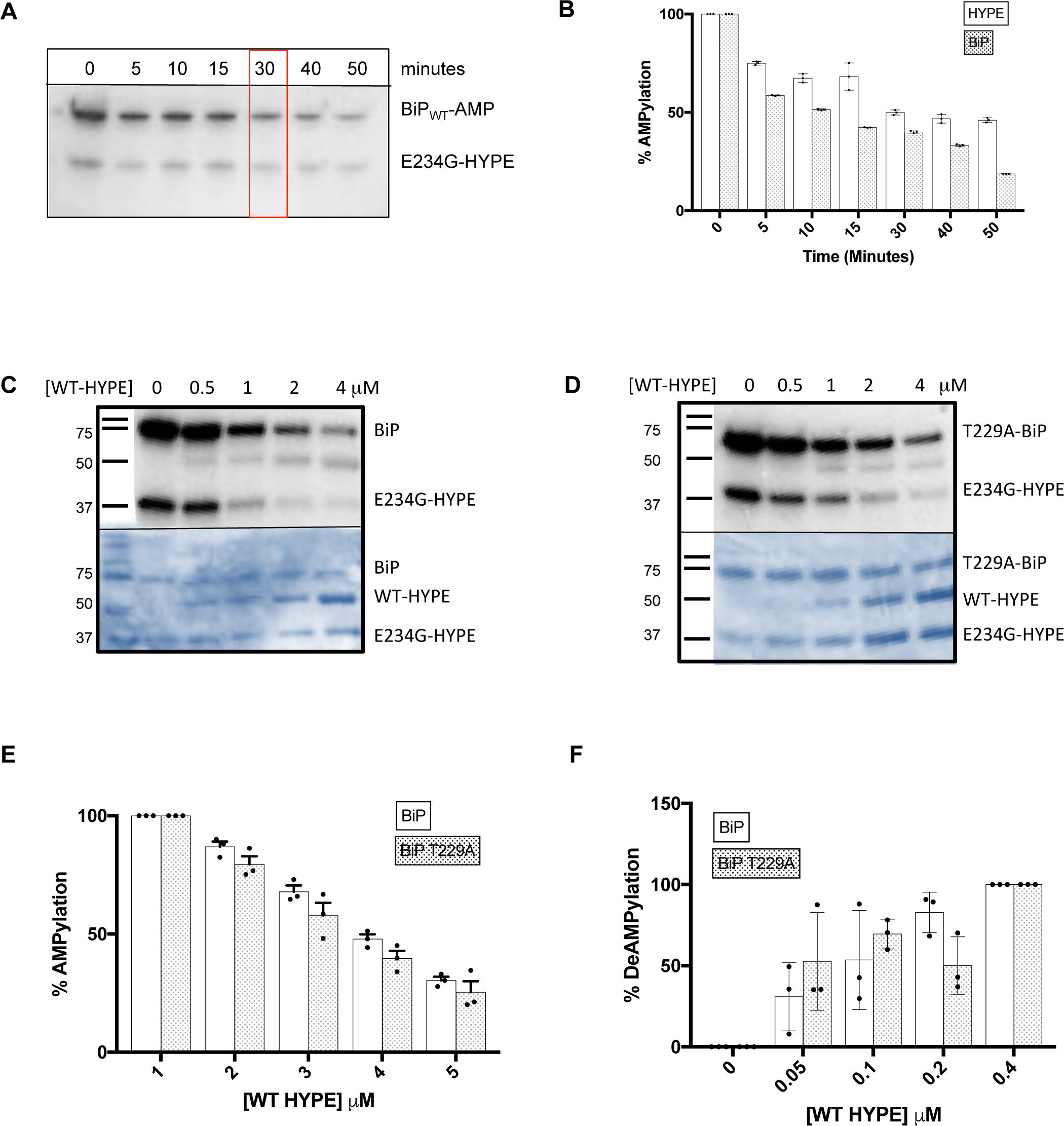
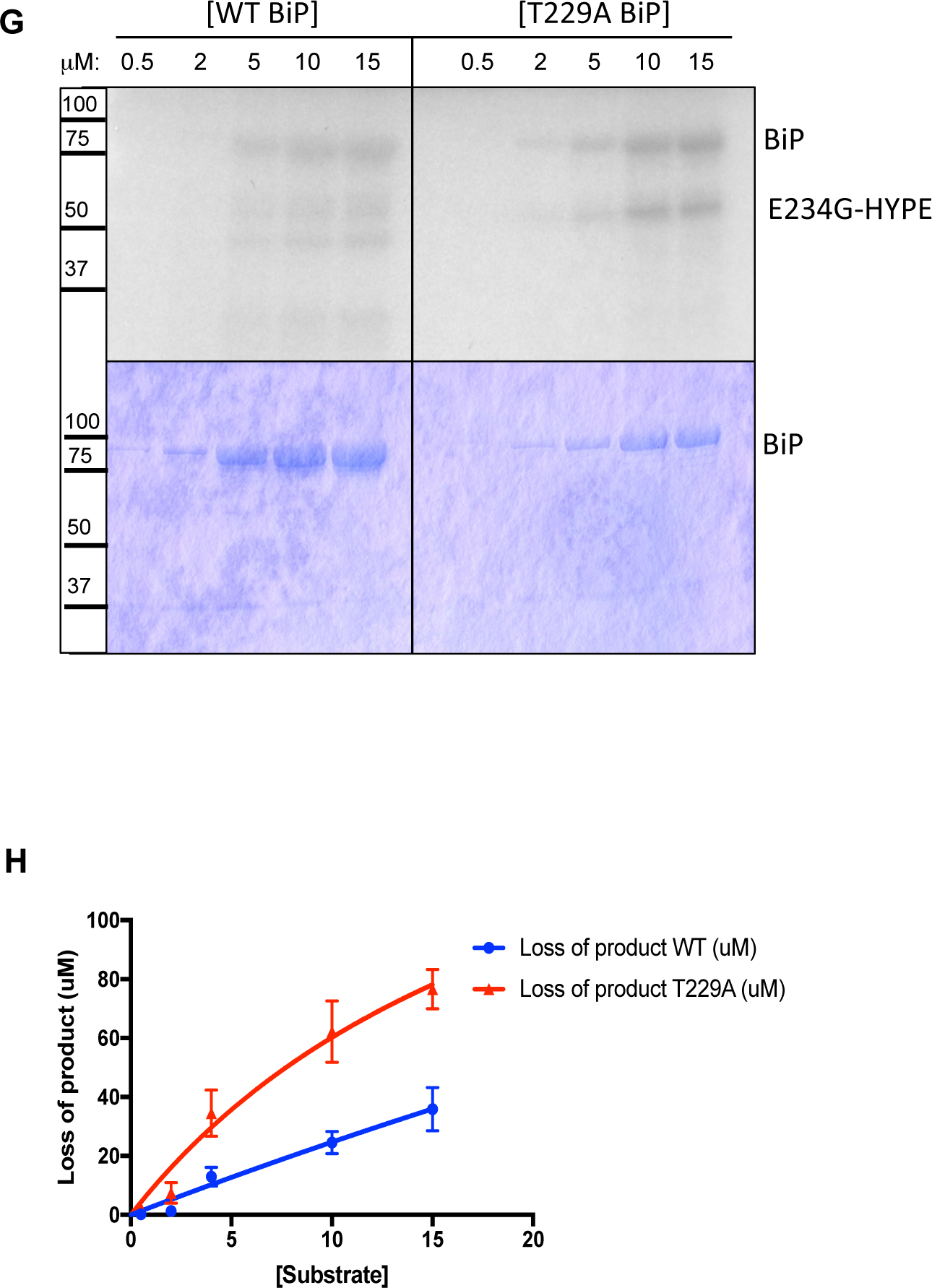
De-AMPylation of adenylylated BiP by WT-HYPE. (A) Determination of the linear range of the de-AMPylation reaction using purified WT-BiP first adenylylated by E234G-HYPE (BiP_WT_-AMP) and WT-HYPE for reaction times varying from 0 to 50 minutes. (B) Quantification of the autoradiograph in (A) to verify the linear range of the de-AMPylation reaction. (C-E) The de-AMPylation of BiP_WT_-AMP versus BiP_T229A_-AMP by increasing concentrations of WT-HYPE was assessed at 30 mins by autoradiography (C, D) and quantified (E). The Y-axis represents the % of radiolabel (i.e., % AMPylation) detected by autoradiography. (F) De-AMPylation reaction of BiP_WT_-AMP or BiP_T229A_-AMP carried out for 30 minutes with increasing concentrations of WT-HYPE and detected using a scintillation counter as the decrease in radioactive counts associated with the adenylylated BiP following treatment with WT-HYPE and washing. The Y-axis represents the % of radiolabel retained (i.e., % de-AMPylation). (G, H) Steady state kinetic analysis of de-AMPylation of WT- and T229A-BiP by WT-HYPE. Increasing concentrations of BiP_wt_-AMP or BiP_T229A_-AMP were incubated with a constant amount of WT-HYPE and adenylylation levels were determined by quantifying corresponding bands on the autoradiograph (G). Quantified bands were normalized against lane 1 (0 μM WT-HYPE + BiP_wt_-AMP) and lane 6 (0 μM WT-HYPE + BiP_T229A_-AMP) of Figure 4G, and fitted to a Michaelis-Menten equation (H).

We next determined whether WT-HYPE displays variable efficiency for de-AMPylating WT-BiP versus T229A-BiP. Following an *in vitro* adenylylation reaction with WT-BiP or T229A-BiP incubated with E234G-HYPE with α^32^P-ATP as described above, the resultant BiP_wt_-AMP or BiP_T229A_-AMP was incubated with increasing concentrations (0.5 μM to 4 μM) of WT-HYPE for 30 minutes, and the level of BiP de-AMPylation assessed by autoradiography (Figures 4C and 4D). The autoradiographs indicate that WT-HYPE de-AMPylates BiP_wt_-AMP somewhat better than BiP_T229A_-AMP, which was confirmed by quantifying the bands on the autoradiograph, however the difference is not significant (Figures 4E). We also repeated the above assay and measured de-AMPylation as a function of released radioactive counts, as detected by a scintillation counter (Figure 4F). Our data show that consistent with the forward adenylylation reaction, the de-AMPylation reaction also shows a small but insignificant preference for the T229A-BiP ATP-bound form of BiP (Figure 4F). We hypothesize that depending on specific cellular signals, the rate of adenylylation versus de-AMPylation reaction differs, leading to elevated levels of either adenylylated or de-AMPylated BiP. Indeed, it was recently reported that while the HYPE dimer functions primarily as an adenylytransferase, HYPE in its monomeric form functions as a de-AMPylase (30). This finding supports our data showing de-AMPylation of BiP and HYPE by WT-HYPE even in the presence of an equal concentration of E234G-HYPE.

The above experiments represent a potential physiological scenario where HYPE is present in both its adenylylation and de-AMPylation competent states. However, to determine the kinetics with which WT-HYPE de-AMPylates BiP_wt_-AMP versus BiP_T229A_-AMP, we need to understand BiP’s de-AMPylation in the absence of any countering adenylylation events. Therefore, we assessed de-AMPylation by first tethering His-tagged E234G-HYPE on a Co2+ column and then incubating it with His-tagged BiP and α^32^P-ATP in an *in vitro* adenylylation reactions (see Methods). Following adenylylation, the column was washed to remove ATP, thus ensuring no further adenylylation could be carried out by E234G-HYPE. Kinetics were determined using a constant concentration of WT-HYPE to de-AMPylate increasing concentrations of BiP_wt_-AMP or BiP_T229A_-AMP, and de-AMPylation was quantified by autoradiography as a measure of radiolabel associated with BiP (Figures 4G and H). Again, our data show that the rate of de-AMPylation of WT BiP is faster than that of T229A BiP, suggesting a link between structural conformation of BiP and its adenylylation status.

### Differential effects of AMPylation on BiP’s ATPase activity

As mentioned earlier, BiP refolds misfolded proteins in the ER by utilizing ATP as an energy source. Consequently, it has the ability to hydrolyze ATP to ADP and inorganic phosphate (Pi). Previously, using a malachite-green based assay that directly measures released Pi as a measure of absorbance at 650 nm, we reported that adenylylation at human BiP’s S365/T366 enhances BiP’s ATPase activity *in vitro* (14). Subsequently, Preissler *et al.* reported that adenylylation at residue T518 on hamster BiP inhibits its ATPase activity and suggested that adenylylation is a mechanism by which BiP activity is stalled as a measure of misfolded protein (client) load (22). The difference in BiP’s ATPase activity associated with Ser365/Thr366-AMP or Thr518-AMP is only two-fold when compared to WT-BiP. This difference is minimal and could be attributed to the use of His-tagged human BiP versus a bulkier GST-tagged hamster BiP, respectively. Therefore, using directly comparable untagged WT-BiP that best represents the native conformation and its corresponding Ser365/T366A and T518A mutant proteins, we assessed ATPase activity of both WT and adenylylation-site mutants of BiP. Specifically, we incubated WT-BiP, T365A/T366A-BiP or T518A-BiP with E234G-HYPE or its catalytically inactive version, E234G/H363A-HYPE, in the presence of ATP and measured the amount of free phosphate released. Our data indicate that following *in vitro* adenylylation, WT-BiP incubated with E234G-HYPE shows a two-fold increase over WT-BiP that was incubated with E234G/H363A-HYPE, in agreement with our previous report that adenylylation enhances BiP’s ATPase activity (Figure 5A). Interestingly, T518A-BiP also displayed a two-fold increase in ATPase activity following incubation with E234G-HYPE, suggesting that this mutant was likely still adenylylated at Thr366 and therefore elicits phenotypes similar to BiP-AMP (Figure 5A). In contrast, even after incubation with E234G-HYPE, S365A/T366A-BiP shows ATPase activity equivalent to unadenylylated WT-BiP, strengthening our original observation that adenylylation at Thr366 enhances BiP’s ATPase activity *in vitro* (Figure 5A). Control reactions showed that the S365A/T366 or T518 mutations themselves did not alter BiP’s ATPase activity (Figure 5B). It was recently reported that AMPylation specifically inhibits BiP’s J-protein assisted ATPase activity, a finding we confirmed by repeating our ATPase assays in the presence of J-domain of ERdJ6 (Figure 5C). Specifically, comparing the left, center, and right sets of histograms in Figure 5C, we find that the ATPase activity of BiP (white bars) is enhanced in the presence of ERdJ6’s J-domain, as compared to WT BiP alone or in the presence of the catalytically inactive J-domain mutant (J*) (Figure 5C and 23). Adenylylation interferes with BiP’s J-protein assisted ATPase activity, as recently reported (compare dark and light grey bars in Figure 5C; and 23). Consistent with all our data, adenylylation causes a meager increase in BiP’s ATPase activity in the absence of the J-domain (Figure 5C, compare white and dark grey histograms in the left set). Thus, we find that while AMPylation inhibits BiP’s ATPase activity during the chaperone cycle governing folding of misfolded proteins, it mildly enhances BiP’s basal ATPase activity.

**Figure 5:**
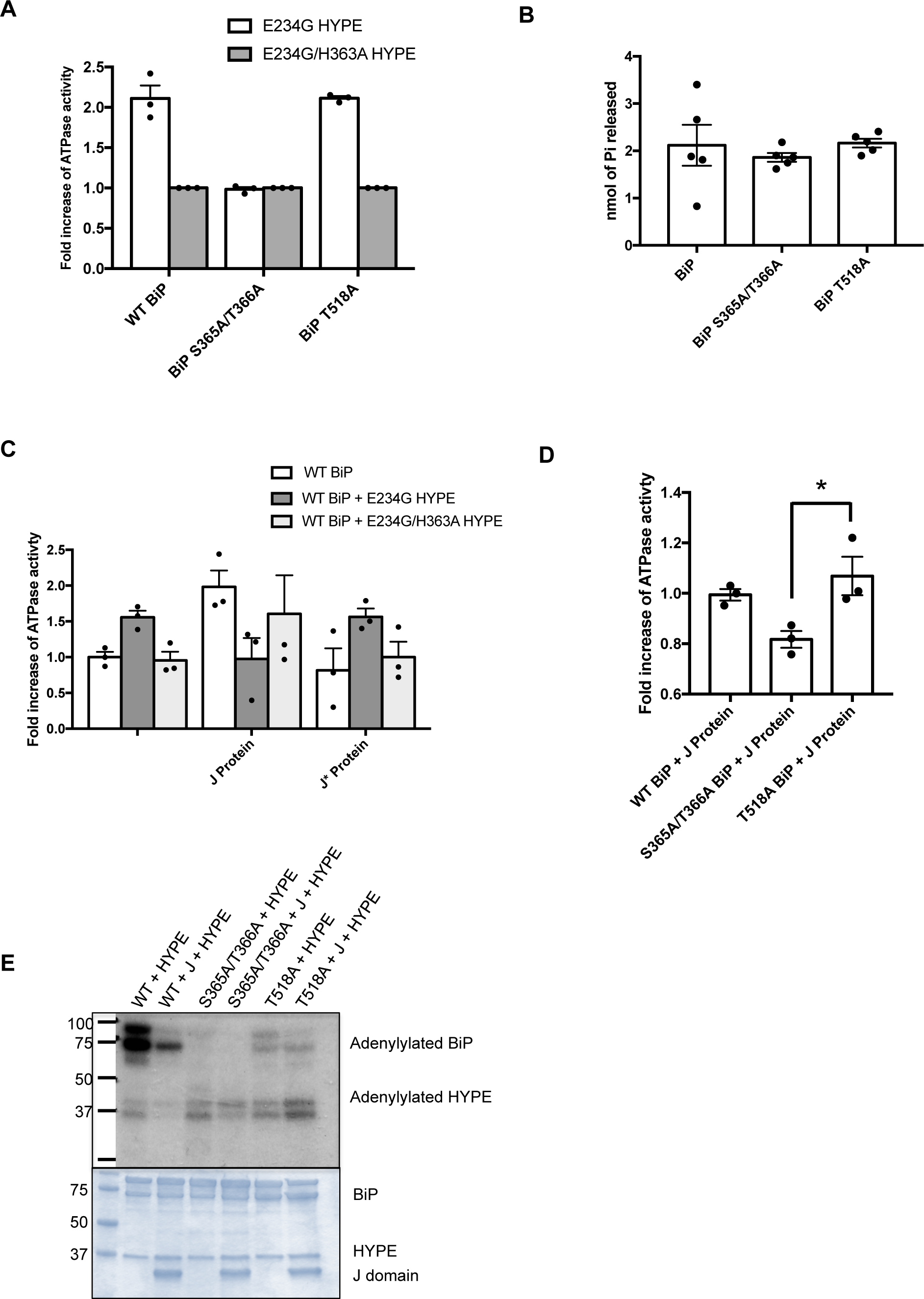
Molecular consequences of BiP adenylylation. (A) WT-, T518A- and S365A/T366A-BiP were assessed for ATPase activity using colorimetric detection of released P_i_ following ATP hydrolysis. Adenylylation was carried out by incubating the BiP substrates with E234G-HYPE prior to the ATPase assay. E234G/H363A-HYPE was used as a negative control for the adenylylation reaction. Adenylylated WT- and T518A-BiP show enhanced ATPase activity when incubated with E234G-HYPE but not with E234G/H363A-HYPE, while S365A/T366A-BiP displays basal level activity irrespective of which HYPE enzyme it is incubated with, indicating a role for adenylylation at the Ser365/Thr366 site. (B) Control ATPase reactions showing that BiP’s S365A/T366A and T518A mutations alone do not alter their ATPase activity when compared to WT-BiP. S365A/T366A and T518A mutants of BiP displayed ATPase activity comparable to WT-BiP, as measured by colorimetric detection of released P_i_. (C) Adenylylation enhances BiP’s basal ATPase activity but inhibits its J-protein assisted ATPase activity. WT-BiP alone or adenylylated by E234G-HYPE or left unmodified by incubating with catalytically inactive E234G/H363A-HYPE was assessed for ATPase activity either alone or in the presence of active ERdJ6 J-protein or catalytically inactive J protein (J*). (D) J-protein assisted ATPase activity of WT-, S365A/T366A-BiP, and T518A-BiP. J-protein assisted ATPase activity of BiP is significantly lower for S365A/T366A-BiP when compared to T518A-BiP. * p = 0.02. (E) Autoradiograph showing adenylylation of WT- or mutant BiP by E234G-HYPE in the presence or absence of J protein.

We also compared the effect of adenylylation on J-protein assisted ATPase activity of WT-BiP and its S365A/T366A and T518A mutants and found that both site mutants displayed ATPase activity at levels similar to WT-BiP (Figure 5D). Interestingly, however, S365A/T366A-BiP’s activity was significantly lower than that of T518A-BiP (*, p = 0.02). We speculate that this augmented difference is because of the opposing activating effects of adenylylation at S365/T366 and inhibitory effects of adenylylation at T518. As a control, we assessed the adenylylation level of each of the proteins in these reactions by autoradiography (Figure 5E). Consistent with our previously published data (14), we find that *in vitro*, majority of the AMPylation specific radiolabel is associated with S365/T366 and that some residual AMPylation, likely on S365/T366, is still observed in the BiP T518 mutant. Interestingly, the cumulative adenylylation on S365A/T366A-BiP and T518A-BiP was less than the adenylylation on WT-BiP, suggesting a possible crosstalk between adenylylation at these two sites. Importantly, we also find that HYPE does not adenylylate the J-domain of ERdJ6 (Figure 5E).

Cumulatively, our data indicate that, in keeping with HYPE’s role in maintaining homeostasis, AMPylation has both activating (basal levels) and inhibitory (J-protein assisted) effects on BiP’s ATPase activity for which both the Thr366 and Thr518 sites are important.

### Structural model for the HYPE-BiP interaction

Similar to other Hsp70 proteins, BiP contains an NBD and an SBD connected by a flexible linker (Figure 1B; {18}). We reported T366, which lies within BiP’s NBD (aa 28-405), as the predominant site of BiP adenylylation by HYPE *in vitro* (14). In agreement, *D. melonagaster* dFic also modified T366, as does *C. elegans* Fic, Fic-1 (21), and a recent report now describes a physiological role for adenylylation at Thr366 of BiP that is important for photoreceptor function and vision in *D. melanogaster* (24). However, using Hamster BiP *in vitro* and in a cellular UPR assay, T518 in BiP’s SBD was suggested to be the only adenylylation site identifiable in cellular assays and was determined to be the only physiologically relevant site (22).

Given the subtle differences in the kinetics of interaction between the various forms of HYPE and BiP, and the debate surrounding the *in vitro* versus *in vivo* significance of the Thr366 and Thr518 sites, understanding how HYPE and BiP interact to form a complex to adenylylate BiP’s different target sites is of paramount importance. Since the crystal structures for WT-HYPE (PDB 4U04), ATP-bound E234G-HYPE (4U07), and ATP-bound T229A-BiP containing both its NBD and SBD (PDB 5E84) are known, we sought to use computational molecular docking to determine how HYPE and BiP interact. By understanding this interaction, we aimed to determine the sites in HYPE and BiP involved in substrate recognition and those involved in mediating adenylylation. Finally, we aimed to identify whether the HYPE-BiP complex structurally favored BiP’s Thr366 in its NBD or Thr518 in its SBD as the site of adenylylation. The HYPE-BiP complex models were generated using a protein–protein docking program, LZerD (32). Based on known characteristics of the structures for HYPE and BiP, we assigned restrictive parameters to our molecular docking analyses to generate the most physiologically relevant HYPE-BiP model. Specifically, since HYPE is predicted to exist as a dimer, we eliminated residues in the HYPE-HYPE dimer interface as points of interaction between HYPE and BiP. Likewise, based on the IbpA-Cdc42 structure, which assigns target recognition to residues outside the Fic enzymatic core, we predicted that residues constituting HYPE’s Fic enzymatic core would also not confer specificity for BiP. Instead, we predicted that HYPE’s TPR domains, which bear similarity to other TPR proteins known to interact with Hsp family proteins [e.g. TPRs of P4H and PP5 proteins bind to Hsp90; (33)], would be responsible for binding to BiP. We also used the ATP-bound E234G-HYPE structure as comparison for ensuring that the Fic active site would accommodate an ATP while interacting with BiP and would not be inhibited by the α_inh_. Finally, we accommodated two orientations for HYPE’s active site, which allow adenylylation of BiP at either Thr366 or at Thr518. Subsequent to our analysis, a structure for adenylylated BiP was reported, which was obtained by filling in a density corresponding to AMP within the structure of a T229A/V461F-BiP mutant which is locked in the same ATP-bound conformation at T229A-BiP (22). Since our molecular docking models were generated using T229A-BiP, the reported BiP-AMP structure did not alter the outcome of our docking computations.

The top scoring resultant models are shown in Figure 6. Figure 6A depicts a model showing a HYPE dimer (in cyan) with BiP (in green) in an orientation that favors adenylylation on Thr366 (in red). This model also shows that the second HYPE binds in an orientation that does not interfere with the bound BiP, suggesting that the second HYPE is likely to bind a second BiP in a similar manner resulting in a 2:2 HYPE-BiP complex (Figure 6A). Interactions between HYPE and BiP occur via HYPE’s TPR domains (in purple) and BiP’s SBD (in light green). The contact sites on BiP where HYPE’s TPR domains interact with BiP’s SBD are shown in dark blue. Specifically, HYPE’s TPR domains interact with residues 554, 555, 557-559, and 561 of BiP’s SBD within a distance of 3.1 Å, allowing hydrogen bonding. Additional interactions occur between four loops (in pink and magenta) at the interface of the Fic active site (in yellow) and BiP’s NBD (in dark green). Figure 6B zooms in at the Fic active site of this model to reveal that Thr366 is positioned at a large distance (17.5 Å) away from HYPE’s active site motif and possibly blocked by the four loops extending each from HYPE (residues 310-325 constituting one loop, in pink) and BiP (residues 47-51, 210-213, and 385-389 constituting three loops, in magenta). However, these loops are unstructured and predicted to be highly flexible relative to the other parts of the protein using FlexPred (34). Therefore, displacement of these loops presents a scenario where Ser365/Thr366 come in close proximity to HYPE’s active site and could be adenylylated.

**Figure 6:**
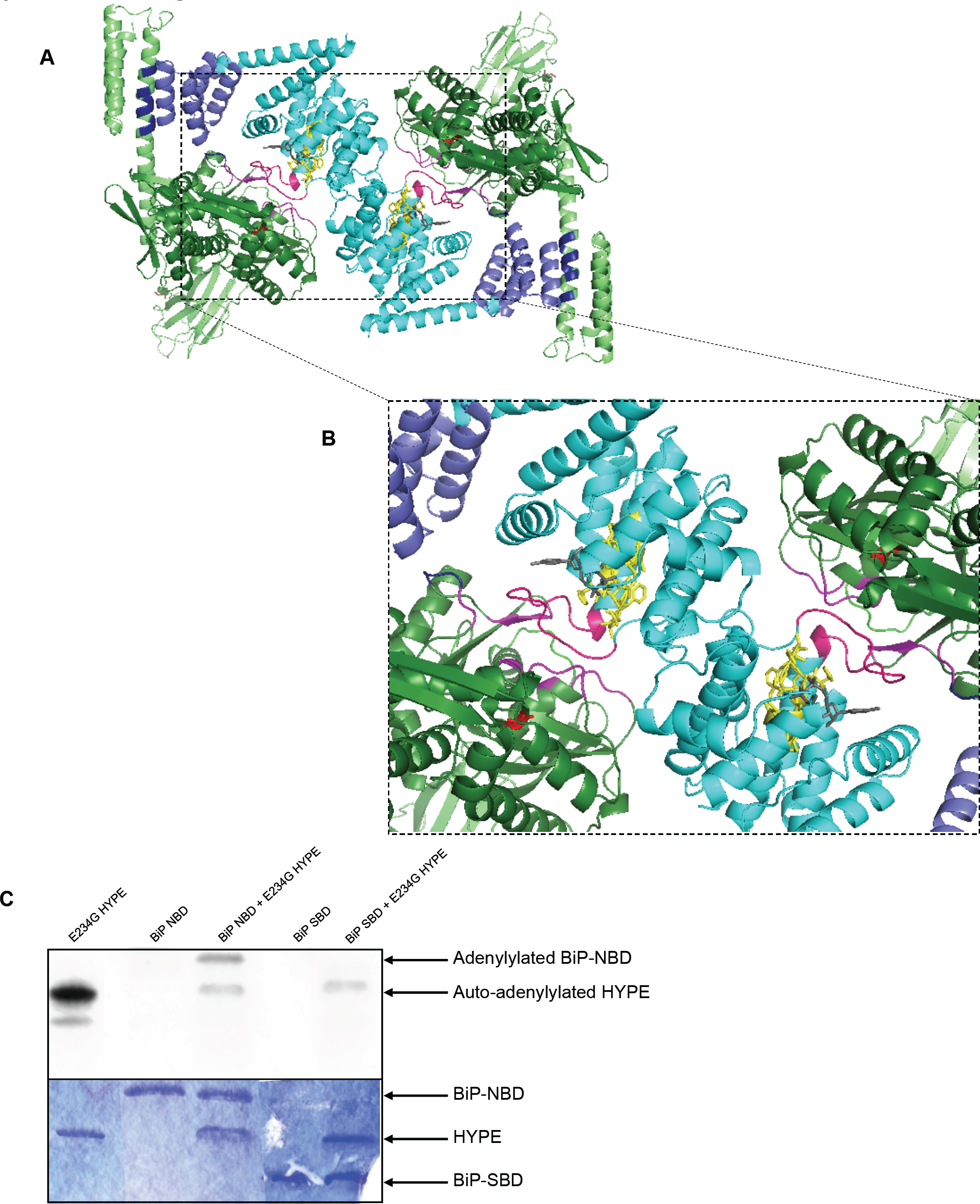
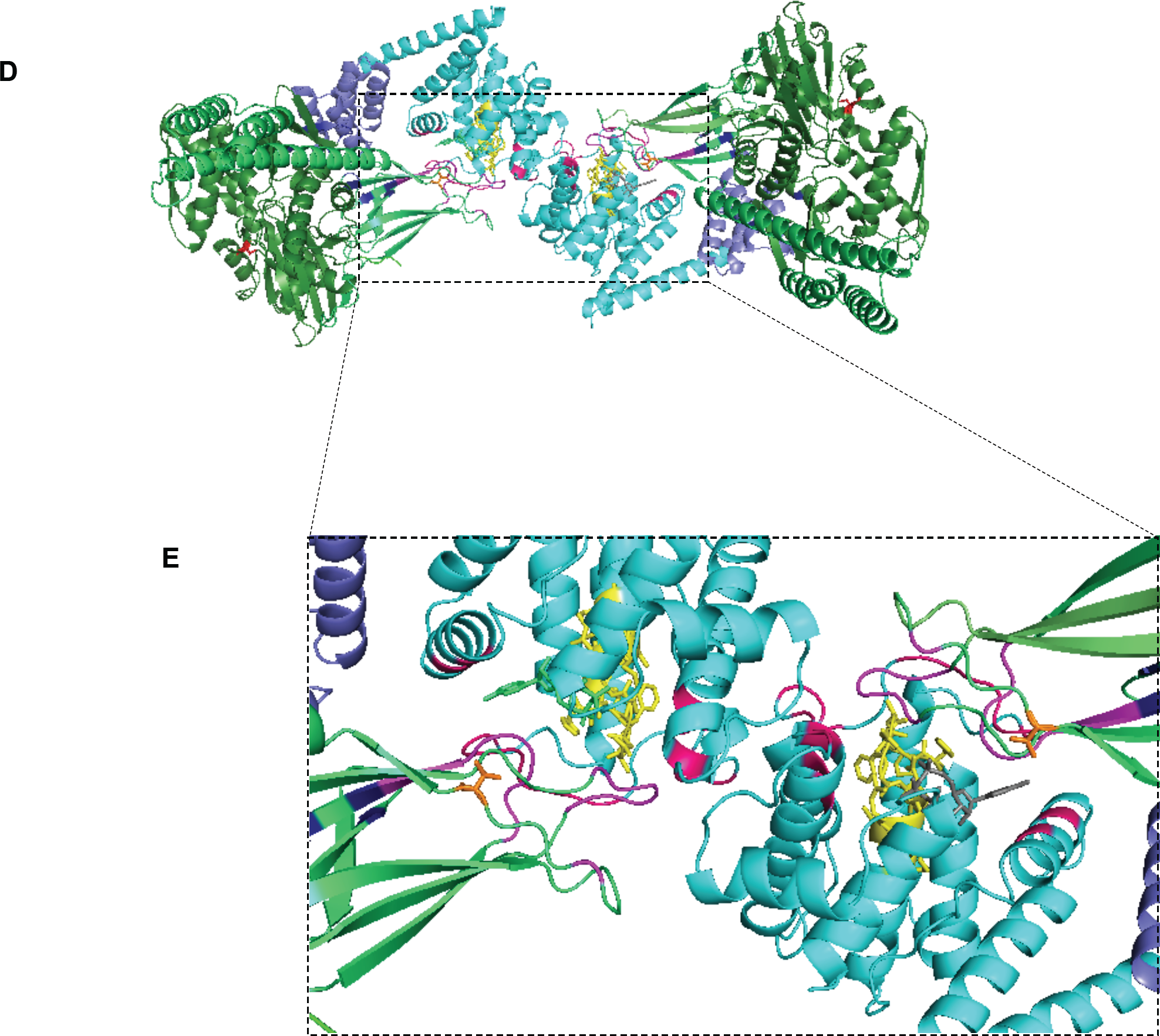
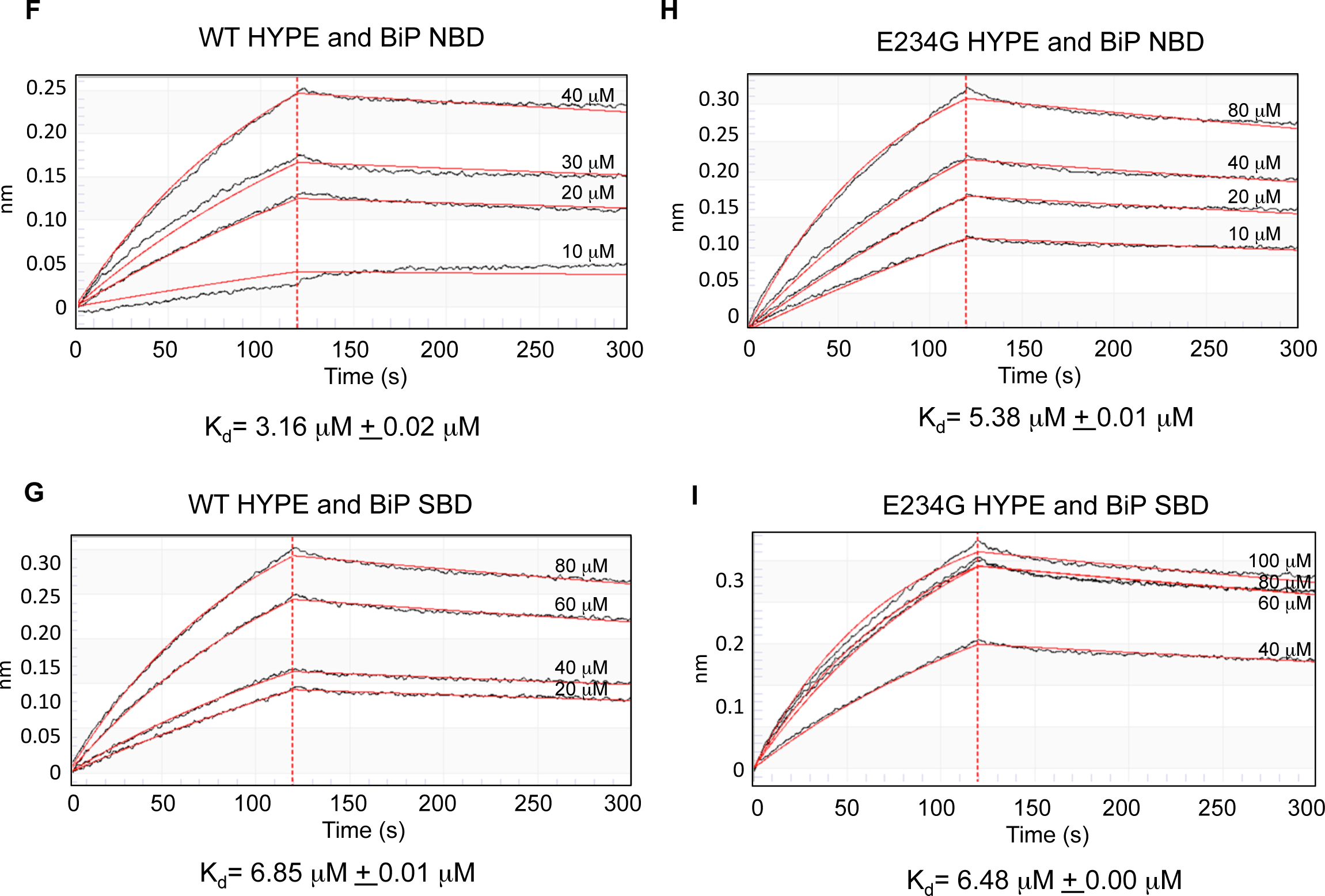
Molecular docking models describing the HYPE-BiP interaction specific for adenylylation at the Thr366 or Thr518 sites. HYPE forms a homodimer via its Fic domain (cyan) and interacts with BiP via its TPR domains (purple). BiP’s NBD (dark green) and SBD (light green) are shown. HYPE’s Fic active site is in yellow, and bound ATP is in grey. Points of contact on BiP between BiP and HYPE at the TPR domain are shown in dark blue, while points of contact around the Fic active site are in magenta. Thr366 is in red. Thr518 is in orange. (A) HYPE dimer bound to two BiP molecules in a conformation that allows adenylylation at Thr366. While this model supports interactions between BiP’s SBD and HYPE’s TPR domains, it also displays extensive interactions between HYPE’s Fic domain (via the pink loop) and BiP’s NBD (via the magenta loops). We predict movement in these flexible loops would allow Thr366 (in red) to gain access to the Fic active site to allow adenylylation at Thr366. (B) Zoomed-in view of the interactions between HYPE and BiP at the Fic active site centered around Thr366 as the adenylylation site. (C) *In vitro* adenylylation assay conducted for 15 minutes with purified BiP-NBD or BiP-SBD and E234G-HYPE shows adenylylation of BiP-NBD and not BiP-SBD, thus validating the model for adenylylation at Thr366. (D) HYPE dimer bound to two BiP molecules in a conformation that allows adenylylation at Thr518. The majority of the docking models support Thr518 as the predominant site of adenylylation. This model supports target recognition via interactions at both BiP’s NBD (with HYPE’s TPRs) and BiP’s SBD (with HYPE’s Fic domain) with adenylylation occurring at Thr518. Thr366 in this model is on an opposite face and would not be amenable to adenylylation in this conformation. (E) Zoomed-in view of the interactions between HYPE and BiP at the Fic active site centered around Thr518 as the adenylylation site. The model shows that Thr518 is not occluded by the flexible loops in HYPE (in pink) or BiP (in magenta), and that these loops show fewer points of contact with each other when compared to their interaction shown in Figure 6B. (F-I) Steady state binding affinity measurements for WT-HYPE or E234G-HYPE and BiP-NBD or BiP-SBD were obtained by Biolayer Interferometry. Constant concentrations of BiP-NBD or BiP-SBD were immobilized on HIS1K sensors and exposed to varying concentrations of WT- or E234G-HYPE. K_d_ values were obtained by fitting the data to a 1:1 global fitting model.

The above model also shows that the four loops at the Fic active site interface allow enough points of interaction between HYPE and BiP to allow the two proteins to interact even in the absence of the BiP SBD. If so, then our model predicts that HYPE should recognize and adenylylate BiP_NBD_ without the need for BiP_SBD_. We, therefore, purified BiP_NBD_ and BiP_SBD_ separately and tested them as targets for HYPE-mediated adenylylation in an *in vitro* reaction using α^32^P-ATP. We found that only BiP_NBD_ is efficiently adenylylated *in vitro* (Figure 6C), thus validating our model in support of Ser365/Thr366 as the adenylylation site.

Figure 6D depicts a model showing HYPE bound to BiP in an orientation that favors adenylylation at Thr518. Figure 6D also reveals that HYPE’s TPR domains interact with two loops in BiP’s NBD - specifically, residues 101-108 constituting the first loop and residues 119-123 constituting the second loop - at a distance of 4.1 Å, which would allow hydrogen bonding between these residues of HYPE and BiP. As in the previous model, binding of BiP does not hamper the HYPE dimer interface, again supporting the formation of a 2:2 HYPE-BiP complex. Figure 6E zooms in on the Fic active site centered around BiP’s Thr518 as the site of adenylylation, and reveals additional sites of interaction between HYPE’s active site loop (in pink) and two loops in BiP’s SBD (shown in magenta) made up of residues 451-458 and 484-489 of BiP. Figure 6E also reveals that the Thr518 on BiP (in orange) is spaced at a distance of 15.8 Å from the Fic active site motif, making it a feasible distance to accommodate an ATP (shown in grey) and allow adenylylation. Together, the information in Figures 6D and 6E indicates that in order for HYPE to adenylylate Thr518 of BiP, interactions must occur with both the NBD and SBD of BiP, and explains why HYPE was unable to modify BiP when its SBD alone was used as a target *in vitro* (Figure 6C).

An alternate explanation for why HYPE fails to modify BiP_SBD_ *in vitro* could be due to a weaker binding affinity between HYPE and BiP_SBD_. Therefore, we tested the steady-state binding affinities of BiP_NBD_ and BiP_SBD_ for HYPE using Biolayer Interferometry. Specifically, we immobilized His_6_-BiP_NBD_ or His_6_-BiP_SBD_ on a His-antibody sensor and titrated increasing concentrations of WT- or E234G-HYPE. The K_d_ of WT-HYPE for BiP_NBD_ was determined as 3.16 μM ± 0.02 μM and for BiP_SBD_ was 6.85 μM ± 0.01 μM, suggesting that both BiP NBD and SBD bind WT-HYPE with similar affinities (Figures 6F and 6G). Results for E234G-HYPE also follow the same trend and displayed similar binding affinities for BiP_NBD_ versus BiP_SBD_ (Figures 6H and 6I). These data suggest that differential binding is not a limiting factor for BiP_SBD_ modification and further supports our model showing the need for interactions at both the NBD and SBD for adenylylation at Thr518. We speculate that binding at BiP_NBD_ presents BiP in the correct orientation to allow adenylylation at Thr518.

## Discussion

Maintenance of ER homeostasis via the UPR pathway is a critical aspect of cell fate, for which BiP plays a central role as a sentinel for UPR activation. Until recently, the main mechanism for regulating BiP acitivity was thought to be via a post-translational ADP-ribosylation event that stalled BiP’s ATPase activity until a threshold client load was reached (35). Specifically, BiP was shown to be ADP-ribosylated at Arg residues R470 & R492 as a means to control UPR activation (35). However, when we reported that HYPE regulated UPR by adenylylating BiP, it became evident that what had previously been thought to be BiP ADP-ribosylation was in fact HYPE-mediated BiP-adenylylation (22). Thus, adenylylation of BiP by HYPE is a key mechanism for regulating how cells cope with ER stress. Understanding the structural and kinetic parameters that govern how HYPE and BiP interact is critical for designing schemes to manipulate this interaction for potential therapeutics.

Our detailed analyses of the kinetic properties governing the HYPE-BiP interaction offer important insights into the efficient and reversible adenylylation of BiP. Our findings are especially valuable for the development and validation of small molecules designed for the purpose of manipulating HYPE’s two enzymatic activities and its interaction with BiP. We show that although E234G-HYPE functions as an efficient adenylyltransferase, both WT- and E234G-HYPE bind to BiP in its ‘open’ and ‘closed’ conformations with similar affinities. It remains to be determined whether BiP’s binding to a misfolded client protein will alter the kinetics of the HYPE-BiP interaction. We also address the lack of consensus with respect to the site of BiP adenylylation and assess the consequence of BiP-adenylylation in the context of both adenylylation sites, Thr366 and Thr518. Preissler *et.al* used a GST-tag (which is comparatively bulky) to assess hamster BiP. Since human and hamster BiP have >90% identity, this disparity in adenylylation site results may be due to the tag used. Therefore, to avoid interference from the tag, we opted to conduct our studies using His-tagged or untagged human BiP.

Our data reaffirm residue Thr366 in BiP’s NBD as the site most associated with an adenylylation signal *in vitro*, and that adenylylation at Thr366 appears to enhance BiP’s basal ATPase activity *in vitro*. The physiological role of BiP’s basal ATPase activity is currently unknown. Our data did not support the inhibition of BiP’s basal ATPase activity via adenylylation at Thr518 *in vitro*, as has been previously reported (22). We reasoned that since the modification at either site on BiP causes only a two-fold difference in ATPase activity *in vitro*, the *in vivo* consequences on ATPase activity, especially in the presence of J proteins and NEFs (Nucleotide exchange factors) that constitute the complete chaperone cycle for bringing clients to BiP could be more substantial. Indeed, it has been reported that adenylylation specifically inhibits BiP’s ATPase activity as stimulated by J proteins (23). Here, we too corroborate an inhibitory role for adenylylation at BiP’s T518 for its J-protein assisted ATPase function. However, comparison of J-protein assisted ATPase activity of the S365/T66 and T518 mutants to WT-BiP reveal subtle but consistent differences in the levels of inhibition (Figure 5D), suggesting a more nuanced control mechanism that balances the effects of modification at each site. Given that both HYPE and BiP are upregulated during UPR, inhibition of BiP ATPase activity by HYPE will be detrimental to the cell. Accordingly, our studies have shown that HYPE is required for cells to cope with prolonged ER stress (14). Such a model of positive regulation of UPR by HYPE would be supported by adenylylation enhancing BiP’s ATPase activity, thus allowing it to refold proteins better. Alternatively, increased ATPase activity may result in faster but incomplete refolding, thus pushing ER stress to be detrimental, thereby supporting an inhibitory effect of BiP adenylylation. It remains to be determined whether different BiP sites could be adenylylated in response to different physiological signals. Further, global phosphoproteomic studies report T518 to be phosphorylated (31), suggesting that perhaps the level or detection of adenylylation *in vitro* at Thr518 may need stringent interactions between multiple sites on HYPE and BiP. Thus, understanding how HYPE could interact and adenylylate the S365/T366 or T518 site on BiP is critical.

Using molecular docking software, we generate a snapshot for the HYPE-BiP complex, which supports Thr518 as the predominant site for BiP adenylylation. This model also identifies residues involved in substrate recognition. Interestingly, models for both adenylylation sites (Figure 6) reconcile the fact that substrate recognition occurs spatially away from the site of enzyme catalysis. This would explain that any effect of adenylylation and interaction with HYPE could affect both the NBD and SBD of BiP. Further, since Thr366 and Thr518 are located on opposite faces of a BiP molecule, it is unlikely that modifications will occur simultaneously on both sites. We speculate that in the context of the cell, during the chaperone cycle when BiP adopts various conformations, both sites could be accessible for adenylylation at different times, with Thr518 proving structurally more favorable for modification. In agreement, the number of model predictions from our molecular docking experiment supporting Thr518 as the site of adenylylation were higher than those supporting Thr366; specifically, 27 of the 29 highest scoring model iterations favored Thr518 over 2 models in support of Thr366.

Finally, we address the kinetics of BiP de-AMPylation by WT-HYPE. Our data show that the rate of de-AMPylation of WT BiP is faster than that of T229A BiP, suggesting a link between the structural conformation of BiP and its adenylylation status. It is perplexing that while measuring the rate of de-AMPylation, we could not complete our reaction to saturation (Figure 4H). One explanation is competitive inhibition of WT-HYPE by E234G-HYPE resulting in non-functional heterodimers. Still, the lack of preference of WT-HYPE towards the pool of BiP molecules in a closed-conformation (BiP_T229A_-AMP) hints at a mechanism of de-AMPylation that differs from what has been speculated from the crystal structure of adenylylated BiP (23). Specifically, adenylylation is speculated to lock BiP in a closed conformation, a conformation that is mimicked by T229A-BiP. It would therefore be expected that WT-HYPE should preferentially de-AMPylate BiP_T229A_-AMP over BiP_wt_-AMP, which our data show is not the case.

Together, our data provide critical information governing substrate specificity and reaction kinetics for HYPE-mediated adenylylation of BiP, and offer keen insights for the design and development of small molecule interventions of this important cellular interaction.

## Methods

### Cloning, Protein Expression and Purification

HYPE clones were obtained by PCR using human HYPE cDNA (from Origene; www.origene.com) as template. For protein production, HYPE wild-type gene encoding amino acids 102-458 and GRP78 (BiP) wild type gene encoding amino acids 19-637 were cloned as an N-terminal His_6_-SUMO fusion in pSMT3 (Addgene; www.addgene.org). Mutations of E234G, E234G/H363A on HYPE and T229A, T518A, S365A/T366A on BiP were made by site-directed mutagenesis. BiP NBD (aa 28-405) and SBD (aa 418-637) were cloned in an N-terminal His fusion in pET45b (Genscript; www.genscript.com).

Recombinant proteins were expressed in *E. coli* BL21-DE3-RILP (Stratagene) in LB medium containing 50 μg/ml of kanamycin (pSMT3) to a density of A600 = 0.6. Protein expression was induced for 12-16 hours at 18°C with 0.4 mM IPTG. Cells were lysed in lysis buffer (50 mM HEPES pH 8.0, 250 mM NaCl, 5 mM imidazole) containing 1 mM PMSF protease inhibitor and purified using a cobalt resin. Resin was washed with wash buffer (50 mM HEPES pH 8.0, 250 mM NaCl, 20 mM imidazole). His_6_-SUMO tag was cleaved by incubating proteins with ULP1 at 4°C. The protein mixture was diluted in wash buffer without imidazole and re-applied to a cobalt column, and flow through containing HYPE was further purified by ion exchange chromatography using 100 mM Tris-HCl pH 7.5 with a salt gradient from 10 mM-1M NaCl. Fractions containing HYPE were purified by size exclusion chromatography in buffer containing 100 mM Tris-HCl pH 7.5, 100 mM NaCl. Protein concentrations were measured using the Bradford method. Purity was determined by SDS-PAGE and proteins were stored at - 80°C. Additionally, each bacterially expressed and purified protein was assessed by mass spectrometry and western blot analysis using antibody raised against AMPylated Tyr (Catalog #ABS184, EMD Millipore) and AMPylated Thr (Catalog #09890, EMD Millipore) to confirm these proteins were not post-translationally modified during the purification process.

### In Vitro Adenylylation Assays

1 μg of wild type or mutant HYPE (aa 102-458) protein was incubated either alone or with 5 μg of BiP NBD or SBD or 5 μg BiP (WT, S365/T366 or T518) or 5 μg J-domain of ERdJ6 in an adenylylation reaction containing 5 mM HEPES pH 7.5, 1 mM manganese chloride tetrahydrate, 0.5 mM EDTA and 0.01 mCi α^32^P-ATP for 15 minutes. Reaction products were separated on 10% polyacrylamide gels and visualized by autoradiography. Bands were quantified using Typhoon phosphoimager and ImageJ software.

### Kinetic Analyses

The adenylylation of BiP by WT- or E234G-HYPE was assayed using [α^32^P] ATP in a reaction buffer containing 5 mM HEPES pH 7.5, 1 mM manganese chloride tetrahydrate, 0.5 mM EDTA, 1 mM ATP and 0.01 mCi α^32^P-ATP in triplicate for 4 minutes. The reaction was stopped with an equal volume of stop solution (0.1 M EDTA, 0.1 M ATP). 25 μl of the stopped reaction were pipetted onto P81 Whatman filter paper. The filters were washed four times in 0.4% phosphoric acid for 30 min, followed by a final wash in 95% ethanol. The filters were then allowed to air dry before being placed in scintillation vials with scintillation liquid followed by counting in a Beckman LS 6000IC scintillation counter. To account for any interfering signals from auto-adenylylation of HYPE, E234G-HYPE was first adenylylated to saturation with 1mM cold ATP before incubating with BiP and α^32^P-ATP.

The rate of the reaction was calculated from a standard curve plotted for DPM versus [ATP]. To determine the linear range of the reaction, constant concentrations of enzyme, substrate and ATP were incubated for different reaction times, the reactions stopped with SDS loading dye, separated on SDS PAGE and visualized by autoradiography. Bands were quantified using Typhoon phosphoimager and ImageJ software and plotted against time. To analyze the apparent kinetic parameters, the ATP and enzyme concentrations were held constant as indicated in the Results while varying the substrate concentration. The data was fitted to the Michaelis-Menten equation using GraphPad Prism.

For the de-AMPylation of BiP, 15 μM BiP was incubated with 1 μM E234G-HYPE in the presence of 1 mM cold ATP and 0.1 mCi α^32^P-ATP for 15 minutes to saturate the adenylylation reaction, followed by addition of increasing amounts (0.5-4 μM) of WT-HYPE. For assessing the kinetics of de-AMPylation, 30 μM His_6_-SUMO BiP bound to a cobalt resin was incubated with 3 μM E234G-HYPE in an adenylylation reaction for 15 minutes, and then washed with adenylylation buffer (5 mM HEPES pH 7.5, 1 mM manganese chloride tetrahydrate, 0.5 mM EDTA) to wash any bound or unused ATP. The adenylylated BiP was aliquoted into 0.5, 2, 4, 10 and 15 μM volumes and incubated with 0.1 μM untagged WT-HYPE in a de-AMPylation reaction for 30 minutes. The reactions were stopped with SDS loading dye, separated on SDS PAGE and visualized by autoradiography. BiP adenylylation was determined using Typhoon phosphoimager and Image J software to quantify the radiolabel associated with BiP bands on the autoradiograph. Loss of adenylylation was quantified by normalizing this BiP-associated signal to the corresponding WT-BiP or T229A-BiP bands incubated with 0 μM WT-HYPE.

### Biolayer Interferometry

The binding kinetic assays were performed on an Octet Red 384 instrument (ForteBio, Menlo Park, CA) with the Biolayer Interferometry (BLI) technique. The BLI technique monitors binding by observing the interference patterns for all wavelength of light in real time, which responds to surface changes at the tip of sensors. The assays were carried out using Anti-Penta-His (HIS1K) biosensors (ForteBio, Menlo Park, CA) at 30°C with the plate shaking speed at 1000 rpm. HIS1K sensors were dipped into the wells of 20 mM His_6-_WT-BiP or His_6_-BiP mutants in HEPES buffer (50 mM HEPES, 150 mM NaCl, pH7.5) for 2 minutes to load BiP on sensors, followed by incubation in buffer for a minute to build a baseline. Then the sensors were dipped into wells of WT- or E234G-HYPE in HEPES buffer at concentrations from 1 μM to 150 μM for 2 minutes to obtain association curves. In the last step, the sensors were dipped back to the baseline HEPES buffer wells for 3 minutes to allow for dissociation curves. Where mentioned, 1 mM ATP was added in the wells containing WT- or E234G-HYPE. The data was analyzed by the ForteBio data analysis software v.9.0.0.12. The binding constants (K_d_) were determined by globally fitting data to the 1:1 fitting model.

### ATPase Assay

ATPase assay was conducted using Abcam Phosphate Assay Kit (ab65622), which measures the abundance of free inorganic phosphate (Pi) released from ATP hydrolysis reaction, that forms a chromogenic complex with malachite green and ammonium molybdate and gives an absorption peak at 650 nm. 1 μM of wild-type, E234G or E234G/H363A-HYPE and 10 μM of BiP and BiP mutants were incubated in an adenylylation reaction under kinetic parameters. Subsequently unmodified or adenylylated BiP was incubated with 1 mM ATP in a 50 μl volume for 30 minutes for the ATPase reaction to occur. The corresponding adenylylation reactions were used as blank samples. The samples were diluted to a final volume of 200 μl in water, 30 μl of phosphate reagent was added and incubated for 30 minutes. Wherever mentioned, 10 μM J-domain of WT (J) or catalytically dead (J*) ERdJ6 was added to the ATPase reaction (23). The absorbance was read at 650 nm using a Tecan spectrophotometer. Phosphate concentration was calculated from a standard curve and represented as a bar chart.

### Molecular Docking

The HYPE-BiP complex models were generated using the LZerD protein–protein docking program (32). LZerD uses a mathematical series expansion of a 3D function called 3D Zernike descriptors (3DZD) to represent protein surfaces (36). This soft representation of protein surface makes the method more tolerant to conformational changes of proteins upon docking. The performance of LZerD was validated by recent rounds of the Critical Assessment of Prediction of Interactions (CAPRI), a community-wide assessment of state-of-the-art docking method (37). Structures deposited in PDB for WT-HYPE (PDB 4U04), and T229A-BiP (PDB 5E84) were used for generating the docking models. The generated docking models were first filtered to exclude models where the HYPE active site was more than 20 Å from both Thr366 and Thr518. The models were further filtered to exclude models where HYPE’s TPR domain was more than 5 Å from BiP. The models were then ranked by the sum of three protein-docking score ranks (37). The top 100 models by this ranking were then filtered to exclude models where the HYPE active site was occluded.

## Acknowledgments

We thank Dr. Clint Chapple for advice on the design of kinetic experiments. We also thank the Purdue University office of Radiological and Environmental Management (REM) and Dr. Nicholas Carpita for access to the Liquid Scintillation Counter. Wild type HYPE_Δ102_/pSMT3 was a gift from Dr. Junyu Xiao and ERdJ6/pGEX-4T1 was kindly provided by Dr. David Ron. We also thank Dr. Linda Hendershot for additional J protein constructs and helpful discussions. Finally, we are grateful to members of the Mattoo lab for their helpful discussions.

## Author contributions

S.M. and A.S. conceived and designed the study. A.S., E.A.Z., B.G.W., and C.C. conducted the experiments. J.M. trained and provided advice on the BLI experiments. D.K. supervised the molecular docking studies. S.M. supervised the overall study. S.M. and A.S. wrote the manuscript. All authors revised and agreed to the final version of the manuscript.

## Abbreviations List

Fic: Filamentation induced by cAMP
AMP: Adenosine monophosphate
HYPE: Huntingtin yeast interacting protein E
UPR: Unfolded protein response
ER: Endoplasmic reticulum
GS: Glutamine synthetase
GS-ATase: Glutamine synthetase adenylyltranferase
BiP: Binding immunoglobulin protein
NBD: Nucleotide binding domain
SBD: Substrate binding domain

